# In vitro Antibacterial effect of *Commiphora myrrha* Oil against Dental Pathogens

**DOI:** 10.1101/2020.10.15.341180

**Authors:** Reem Izzeldien, Sondos Abdulaziz, Ayat Ahmed, Mounkaila Noma

## Abstract

Dental caries is a chronic disease caused by the interaction of oral microorganisms, diet and host factors over time. *Streptococcus mutans* is considered as the main bacteria involve in dental decay, while the level of *Lactobacillus spp*. is directly related to the presence or onset of caries. *Commiphora myrrha* is an ancient plant which extracts are used as antiseptic and anti-inflammatory for mouth and throat due to it is antimicrobial activity. This study assessed the impact of *Commiphora myrrha* on two bacteria *Streptococcus mutans* and *Lactobacillus spp*. involved in dental caries. Three samples of *Streptococcus mutans* bacteria were collected randomly from patients with dental caries in Khartoum dental teaching hospital, while *Lactobacillus spp*. were obtained from fermented milk. Disk and well diffusion methods were used to test the effect of four concentration (100,50,25 and 12.5 mg/ml) of Myrrha volatile oil, extracted by hydro-distillation technique. The biochemical analysis of *Commiphora myrrha* oil was carried out using Gas Chromatography/Mass Spectrophotometric technique. The finding revealed that the four concentrations of oil were effective on *Streptococcus mutans* with the largest inhibition zone (18.7± 0.6 mm) through the well diffusion method and inhibition zone of (14.00 mm) with disc diffusion method regardless the two methods these inhibition zones were recorded at 100 mg/ml, with Minimum Bactericidal Concentration (MBC) at 3.125 mg/ml. While *Lactobacillus spp*. bacteria sensitive to three concentrations (100,50 and 25 mg/ml) and resistance to concentration 12.5 mg/ml, it is MBC found to be 25 mg/ml. In conclusion, this research revealed that Myrrh oil is effective on both *S.mutans* and *Lactobacillus spp*. Hence, Myrrha oil is a potential antibacterial product of interest in dental caries.

## Introduction

Dental caries is a chronic disease caused by the interaction of oral microorganism in dental plaque, diet and host factors (including teeth and saliva) over time, this result in localized destructions of hard tissues of teeth. Various factors include social, environmental, genetic, biochemical, and immunologic factors WHO declares that deprived oral health and it is related diseases may have dreadful effect on common health as well as eminence of life. Dental caries is a common and major public health oral disease which hampers the attainment and protection of oral health in different age groups [1,2]. Diet type play an important role in caries formation, as fermentable sugar and carbohydrates interact with the acidogenic bacteria in the mouth leading to acid formation which result in tooth decalcification and destruction of hard parts of the tooth [3,4]. *Streptococcus mutans*, one of the acidogenic bacteria, is the main bacteria responsible of dental decay, as indicated by epidemiological studies which revealed that 74% to 100% of the bacteria associated with dental decay was *Streptococcus mutans*; *Lactobacillus* is another acidogenic bacteria which level had a direct relation with the presence or onset of caries [5].

*Streptococcus mutan* is a Gram-positive lactic acid bacterium which produces energy by glycolysis and metabolizes large amount of carbohydrates which is its characteristic cariogencity signature. *S.mutans* lives naturally in dental plaque with several other microorganisms; but the ability of *S.mutans* bacteria to adapt in environmental change is the key reason explaining that the bacteria is a major etiological agent of dental caries. Furthermore, it has the ability to synthesize large amount of extracellular polymer of glucan from sucrose. Strains of *mutans* bacteria are classified based on the composition of cell-surface rhamose-glucose polysaccharide in four serological groups (c, e, f, k); it was reported 75 % of the bacteria on dental plaque are from group c [6]. *Lactobacillus spp*. is an opportunistic Gram-positive bacilli bacteria that needs specific environmental requirement to sustain. This includes access to fermentable carbohydrate, anaerobic niches and low Ph environment as mostly found in dental lesion, stomach and vagina. This last is considered as one of the ways of transmitting the bacteria from mother to the child during birth. Another source of *lactobacilli* transmission is contaminated food and infected human [7].

Some plants are used since the Bible time for wound dressing, women health and beauty care. *Commiphora myrrha* is one of this ancient plant used as antiseptic and anti-inflammatory for mouth and throat. It is from genus *Commiphora* and *Burseraceae* family, which are found in southern Arabia, Somalia, Kenya and some Asian countries. Myrrh composed of alcohol-soluble resin, water-soluble gum which contain polysaccharides and proteins, and volatile oil that composed of steroids, sterols and terpenes [9]. Myrrh volatile oil is extracted by distillation method (hydro-distillation). Furano-type compounds has been reported as major constituents; other components were also reported furanodiene, 19.7%, furanoeudesma-1,3-diene, 34.0%; and lindestrene, 12.0% as the major constituents of Ethiopian species [10]. Herbal extract is used in dental care as tooth cleansing, anti-inflammatory, antimicrobial agent, and analgesic. WHO stated that 80% of the world depends on herbal medications for their primary health care because of the safety and the low cost. *Commiphora myrrha* is used in dentistry for malodour and as anti-inflammatory in periodontitis as it promotes healing [11]. Twenty-three types of 100% natural essential oils were tested to assess their antimicrobial effects on oral strain using disk diffusion method in Korea [12]; The findings indicated that seventeen oils were effective on *S.mutans*, with myrrh, basil, and carrot seed which had a high antimicrobial activity. Of those 17 oils, myrrh had the highest antimicrobial activity with inhibition zone 18.34 mm±0.26. Eighteen oils were found effective against *Porphyromonas gingivalis*; the high antimicrobial activity was obtained from tea tree, carrot seed, and cinnamons. Sixteen oils had high antimicrobial activity on *Lactobacillus rhamnosus*, these oils were from carrot seed and peppermint cinnamon. The antibacterial efficacy of Chlorohexidine Gluconate (0.2%) and Saudi Myrrh (1g % w/v aqueous colloidal solution) mouthwashes were compared on oral microbial flora and other microorganisms on a sample of female participants. The nine microorganisms tested were *Escherichia coli* 25922, *Salmonella* 25566, *Klebsiella pneumonia* 13883, *Pseudomonas* 27853, *Proteus, Staphylococcus aureus* 25923, *Streptococcus pneumonia, Streptococcus mutans*, and *Candida albicans*. The results of the test revealed that Myrrh produced antimicrobial activity against *S.aureus, S.mutans* and *Candida albicans* and other microorganisms that comparable to Chlorohexidine gluconate [13]. The pharmaceutical formulations of mouthwashes, containing extracted Yemeni Myrrh as a single active constituent indicated that the hydro-alcohol extracted by ethanol (phosphate buffer pH 7 (85:15)) had an antimicrobial activity. Of 10 pharmaceutical formulations of Myrrh tinctures prepared and tested, two formulations, containing 9.5% (M9) and 10.5 % w/v (M10) of sodium lauryl sulphate had satisfactory results. In addition, the antimicrobial activity of the formulation M9 was higher than those of two commercial mouthwashes and one oral antifungal suspension [14]. Myrrh has also an antimicrobial activity; It is used for treatment of oral ulcers, gingivitis, sinusitis, glomerulonephritis, brucellosis and a variety of skin disorders [15]. Myrrh has a known anti-inflammatory effect proved by it is ability to supress interleukin 31 (IL-31) mRNA and IL-31 protein expressions, production of IL-31cytokine and inhibits histamine production. Our research aimed to assess the effect of *Commiphora Myrrha* on *Streptococcus mutans*, and *Lactobacillus spp*. bacteria as causes of dental caries and contribute in providing scientific-based evidence of the antibacterial effect of Myrrh oil.

## Materials and methods

An experimental study was done from August 2019 to February 2020.

The swaps were obtained from outpatients of Khartoum dental teaching hospital. Isolates of lactobacillus bacteria were received from Sudan University of Science and Technology. The Laboratory of microbiology department of the Faculty of Medical Laboratory Sciences of the University of Medical Sciences and Technology was used to perform the laboratory tests.

### Preparation and Extraction *of Commiphora myrrha* oil

Plant sample obtained from a local shop in Omdurman market in August 2019. The specimen was deposited in the herbarium of medicinal and aromatic plants institute (MAPTRI), Khartoum, Sudan. The fresh sample was cleaned, air dried and ground to fine powder using a pestle and mortar. Figure 1: Myrrh plant (oleogum resin part)

**Figure 1:**
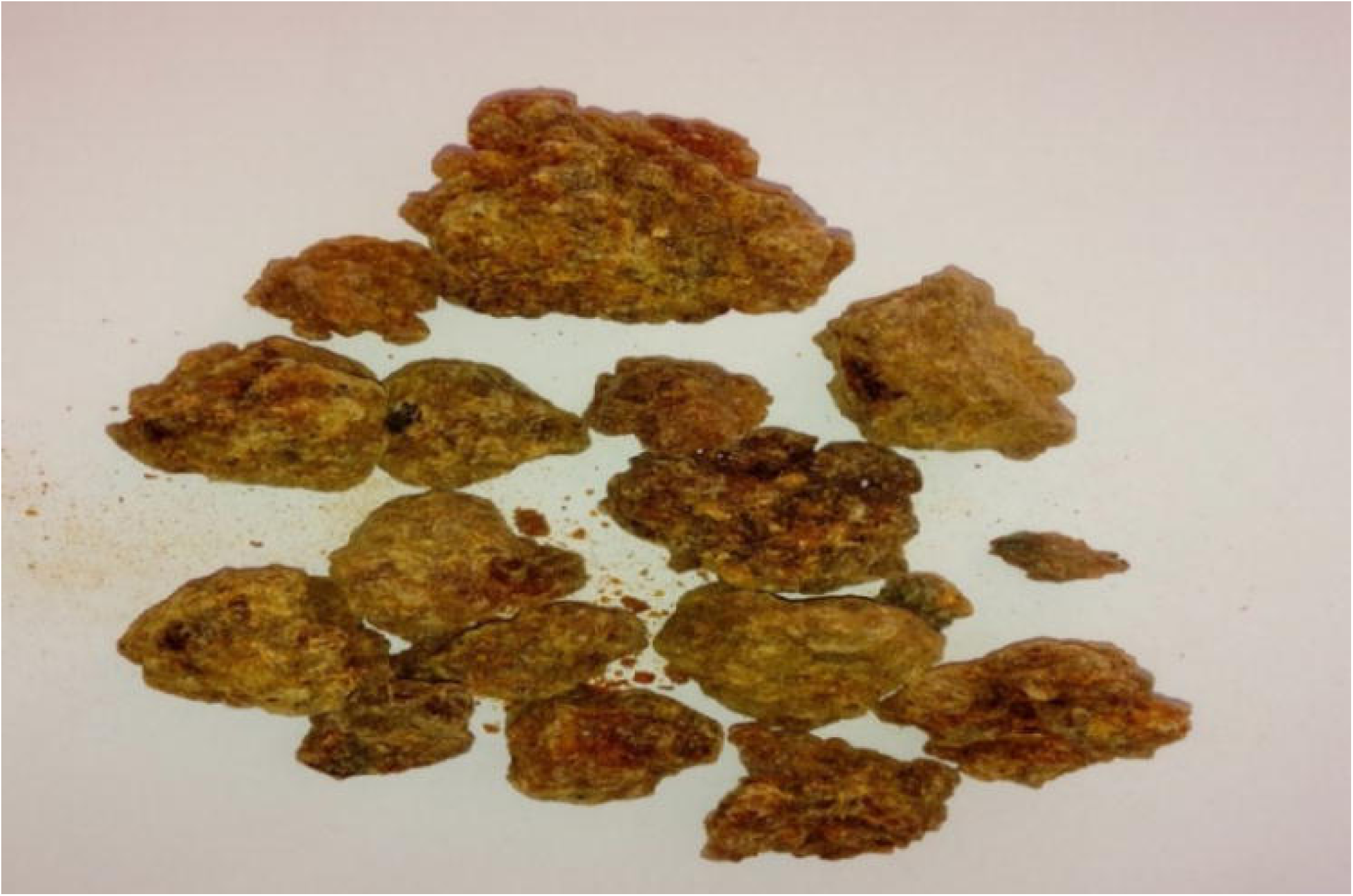
Myrrh plant (oleogum resin part)

About 250 grams of the sample weighted and extracted by hydro-distillation technique [9] this experiment was done in Environmental and Natural Resources and Desertification Research Institute (ENDRI), Khartoum, Sudan. The oil yield was calculated.

### Bacterial strains

#### Lactobacillus spp

bacteria were obtained from Department of Diary Science and Technology of the Faculty of Animal Production, Sudan University of Science and Technology. The bacteria were isolated from traditional fermented milk (berkeep) and the department identified and availed to us three isolates.

#### Streptococcus mutans

isolated from dental caries. Samples were collected randomly in Khartoum Dental Teaching Hospital from patients with dental caries, sterile cotton swabs were used to take the swabs from inside carious teeth, the swabs were transported in Amies transport medium till delivery to the laboratory.

### Identification of S.mutans bacteria

The caries swabs were inoculated on sterile Mitis Salivaries Agar plates. The plates incubated anaerobically at 37□ overnight, microscopic identification made then subculture in Muller Hinton Agar supplemented in blood plates, for further identification biochemical test applied [16].

### Microscopical examination

Dried and fixed smear were prepared from culture media. Gram stain were applied crystal violet stain for 1 minute, washed with tape water and decolorized by alcohol for few second, washed immediately with tape water and covered with safranin for 2 minutes then washed again, allowed to dry and examined microscopically using oil immersion lens (X100).

### Biochemical tests for identification of isolates

#### Antibiotic sensitivity test

Optochin discs were used to test sensitivity of isolates [16].

#### Sugar fermentation tests

Sugar fermentation tests and Esculin hydrolysis are commonly tests used for identification and confirmation of bacteria [17].

#### Phenol red indicator preparation

About 0.25 gram of phenol red powder added to 10 ml of methanol and 10 ml distilled water and mixed.

#### Sucrose

About 1 gram of sucrose sugar was dissolved in 100 ml of peptone water, then 1 ml of phenol red was added to sugar. The bacteria were put to test tube from the media and incubated for 24 h at 37 □ then the colour changed were observed.

#### Glucose

About 1 gram of Glucose sugar was dissolved in 100 ml of peptone water, then 1 ml of phenol red was added to sugar. The bacteria were put to test tube from the media and incubated for 24 h at 37 □ then the colour changed was observed.

#### Mannitol

About 1 gram of Mannitol sugar was dissolved in 100 ml of peptone water, then 1 ml of phenol red was added to sugar. The bacteria were put to test tube from the media and incubated for 24 h at 37 □ then the colour changed was observed.

#### Lactose

About 1 gram of Lactose sugar was dissolved in 100 ml of peptone water, then 1 ml of phenol red was added to sugar. The bacteria were put to test tube from the media and incubated for 24 h at 37 □ then the colour changed was observed.

If colour changed to yellow = positive results

If colour remain red = negative results

#### Esculin test

The surface of aesculin agar bacteria inoculated using sterile loop and incubated at 37 °C for 24 h development of black colour indicates positive result.

#### Voges Proskauer (VP)test

Bacteria inoculated in Methyl red-Voges-Proskauer broth and incubated for 24 h at 37°C, after that 1 ml of 40% KOH and 3 ml of 5% solution of α-naphthol were added. A positive reaction indicated by development of eosin-pink colour.

#### Catalase test

Test used to differentiate bacteria produce enzyme catalase such as *Staphylococci* from non-catalase producing such as *Streptococci*. In sterile glass slide drops of hydrogen peroxide H_2_O_2_ added, using sterile wooden stick several colonies were removed and immersed in, then immediate bubble production observed.

### Antimicrobial activity of Myrrh oil

Two different methods were used to detect antibacterial activity of Myrrh oil; disc diffusion method and well diffusion method.

#### Disc diffusion method

The antibacterial activity of plant extract was determined using the disc diffusion method [17]. Twenty ml aliquots of Mueller Hinton agar will distributed into sterile Petri-dishes. About 0.1 ml of the isolates and standardized bacterial stock suspension 10^8^CFU mL-1 will streaked on Mueller Hinton agar medium plates using sterile cotton swab. Standardized sterilized filter paper discs 6 mm (What man NO1) diameter will soaked in the prepared extracts, then were placed on surface of the test bacteria plates. The plates incubated anaerobically at 37□ for 24 hour and the zone of inhibition (mm) was measured.

#### Well diffusion method

The second antibacterial activity of plant extract was determined using Well diffusion method [18].

#### Determination of minimum bactericidal concentration (MBC)

The lowest concentration of each antimicrobial agent that inhibits the growth of the microorganisms being tested is known as minimum inhibitory concentration (MIC) and is detected by lack of turbidity matching with a negative control. Furthermore, the minimum bactericidal concentration (MBC) is defined as the lowest concentration of an agent killing the majority of bacterial inoculums [19].

To determine the MBC of *C.myrrha* oil four concentrations were prepared by serial dilution starting by stokes oil concentration (12.5 mg/ml). Each tube seeded with bacteria prepared according to Mcfarland 0.5 turbidity standard and incubated in 37□ for 24 h **“**Fig 2”.

**Figure 2:**
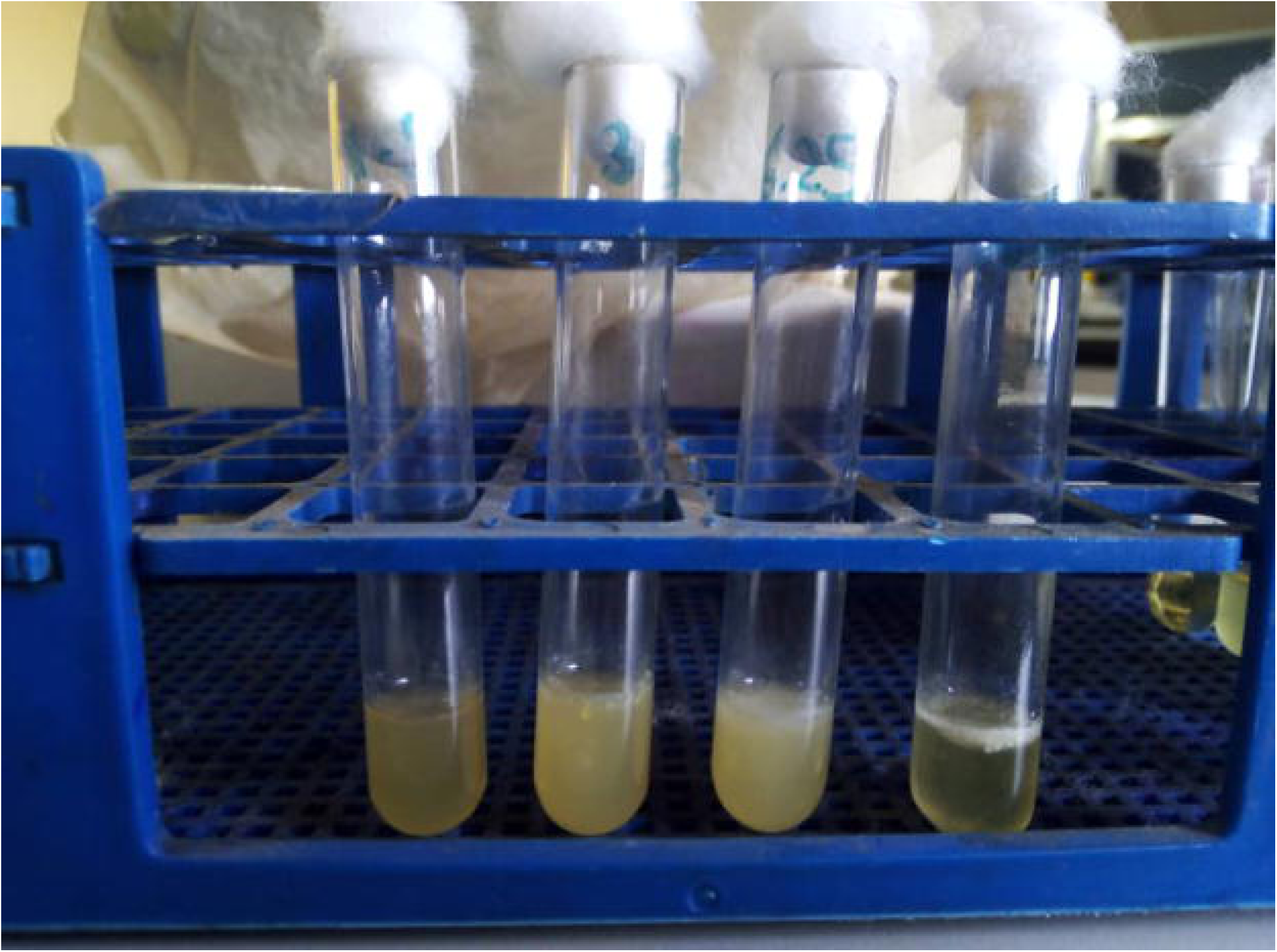
Determination of MBC of Myrrh oil. After 24 h, each tube swabbed in plate of Muller Hinton agar suspended in blood using sterile cotton swab and incubated anaerobically for 24 h in 37□, results were recorded to observe the growth of bacterial colonies.

### Gas Chromatography Mass Spectrometry (GC/MS)

The qualitative and quantitative analysis of the sample was carried out by using GC-MS technique model (GC-MS-QP2010-Ultra) from japans ‘Simadzu Company, with capillary column (Rtx-5ms-30 m×0.25 mm×0.25 µm).The sample was injected by using split mode, Helium as the carrier gas passed with flow rate 1.61 ml/min. The temperature program was started from 60°C with rate 10°C /min to 300°C as final temperature degree with 2 minhold time: the injection port temperature was 300°C. The ion source temperature was 200°C and the interface temperature was 250°C. The sample was analyzed by using scan mode in the range of m/z 40-500 charges to ratio and the total run time was 26 min. Identification of components for the sample was achieved by comparing their retention times and mass fragmentation patterns with those available in the library from the National Institute of Standards and Technology (NIST).

## Results

### Yield percentage of *C.myrrha*

*C.myrrha* volatile oil yield was found to be 0.8% using hydro-distillation method.

### Identification of *S.mutans*

Microscopic examination revealed Gram-positive cocci in chains. That produced α-haemolysis in Muller Hinton agar supplemented with blood and resistant to Optochin disks “Fig 3”

**Figure 3:**
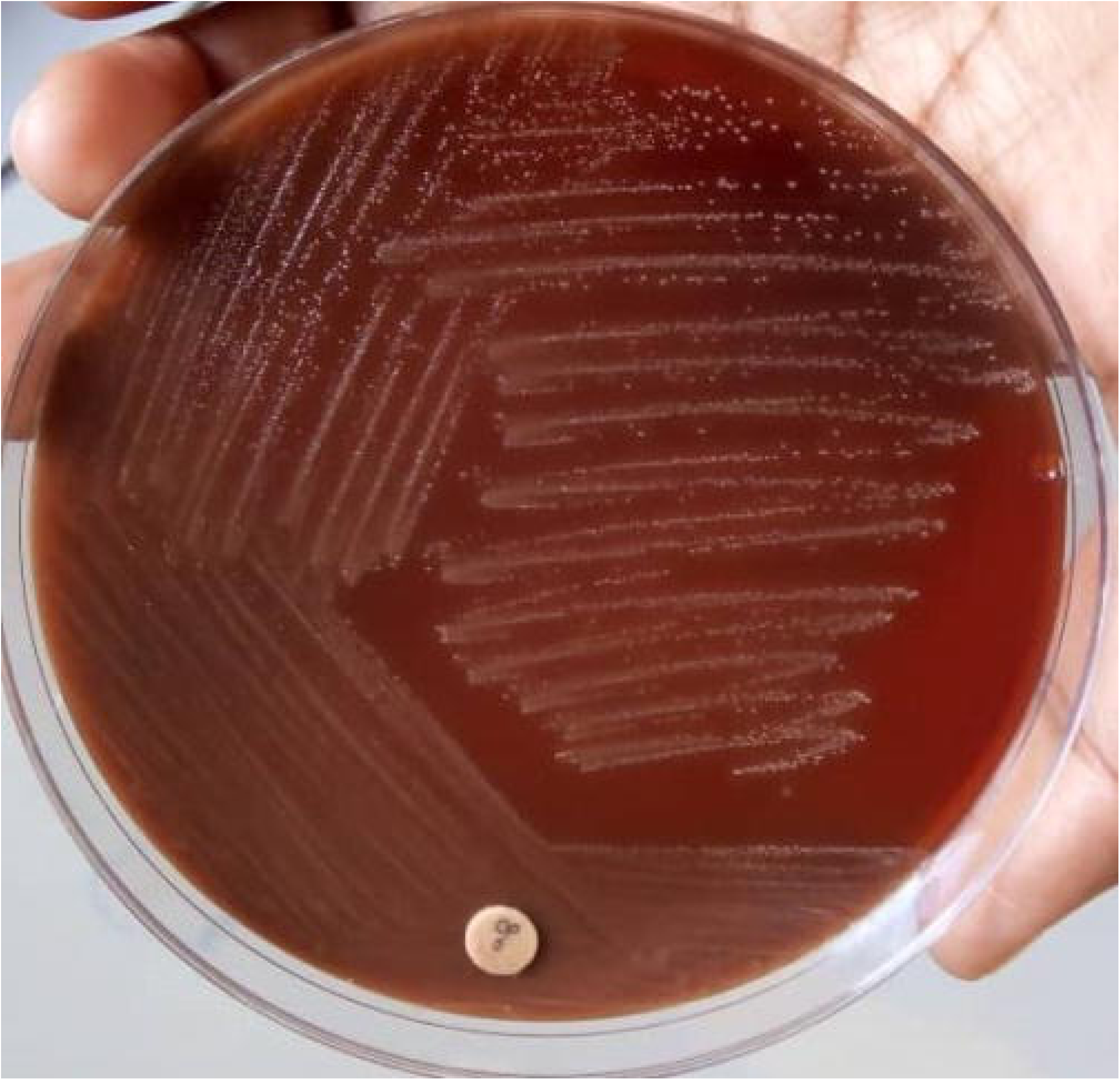
Bacteria with α-haemolysis resistant to optochin disk in Blood agar. All 3 isolates fermented the four types of sugars (Glucose, Sucrose, Lactose and Mannitol). the fermentation was detected by the turning of the colour of phenol from red to yellow “Fig 4”. Esculin and V.P tests were positive. the bacteria were catalase negative. As a final result all three isolated bacteria were identified as *Streptococcus mutans* bacteria.

### Antimicrobial activity

Disk and well diffusion methods were used to test the effect of four concentration (100,50,25 and 12.5 mg/ml) of Myrrha volatile oil against Six bacteria strains. Disc diffusion method represented 66.7% (6/9) of the method and the remaining 33,3% (3/9) was Well method as revealed by “**Table 1**”

**Table 1:** Types of bacteria (*Streptococcus mutans* and *Lactobacillus*) and testing methods used.

| Bacteria | Method of testing |  | Total |
| --- | --- | --- | --- |
|  | Disc diffusion method | Well method |  |
| *Streptococcus mutans* bacteria 1 | 1 | 1 | 2 |
| *Streptococcus mutans* bacteria 2 | 1 | 1 | 2 |
| *Streptococcus mutans* bacteria 3 | 1 | 1 | 2 |
| *Lactobacillus* bacteria 1 | 1 | 0 | 1 |
| *Lactobacillus* bacteria 2 | 1 | 0 | 1 |
| *Lactobacillus* bacteria 3 | 1 | 0 | 1 |
| **Total** | **6** | **3** | **9** |
| % | 66.7 | 33.3 |  |

Inhibition zone of *S,mutans* was measured across four concentrations of oil using two methods as revealed by “**Table 2**”.

**Table 2:** Inhibition zones of *S.mutans* and *Lactobacillus ssp*. across the four concentrations.

| Variables | Myrrh volatile oil concentrations<br>(mg/ml) |  |  |  |
| --- | --- | --- | --- | --- |
|  | 100 | 50 | 25 | 12.50 |
| **Inhibition zone for *Streptococcus mutans* bacteria using disc diffusion method (n=3)** |  |  |  |  |
| Mean | 14.0 | 12.7 | 11.7 | 10.7 |
| Std. Deviation | 0.0 | 0.6 | 0.6 | 0.6 |
| Minimum | 14.0 | 12.0 | 11.0 | 10.0 |
| Maximum | 14.0 | 13.0 | 12.0 | 11.0 |
| **Inhibition zone for *Streptococcus mutans* bacteria using well method(n=3)** |  |  |  |  |
| Mean | 18.7 | 17.7 | 15.7 | 14.0 |
| Std. Deviation | 0.6 | 0.6 | 0.6 | 1.7 |
| Minimum | 18.0 | 17.0 | 15.0 | 12.0 |
| Maximum | 19.0 | 18.0 | 16.0 | 15.0 |
| **Inhibition zone for *Lactobacillus ssp.* using disc diffusion method (n=3)** |  |  |  |  |
| Mean | 10.0 | 8.3 | 6.7 | 0.0 |
| Std. Deviation | 2.0 | 1.5 | 1.2 | 0.0 |
| Minimum | 8.0 | 7.0 | 6.0 | 0.0 |
| Maximum | 12.0 | 10.0 | 8.0 | 0.0 |

In both disc diffusion method and well method the highest inhibition zones were recorded in 100 mg/ml (14.0 mm and 18.7 mm ± 0.6) respectively.

Whereas the highest inhibition zone against *Lactobacillus ssp*. was found to be 10.00 mm ± 2 at 100 mg/ml. “**Fig 5**”

**Figure 4:**
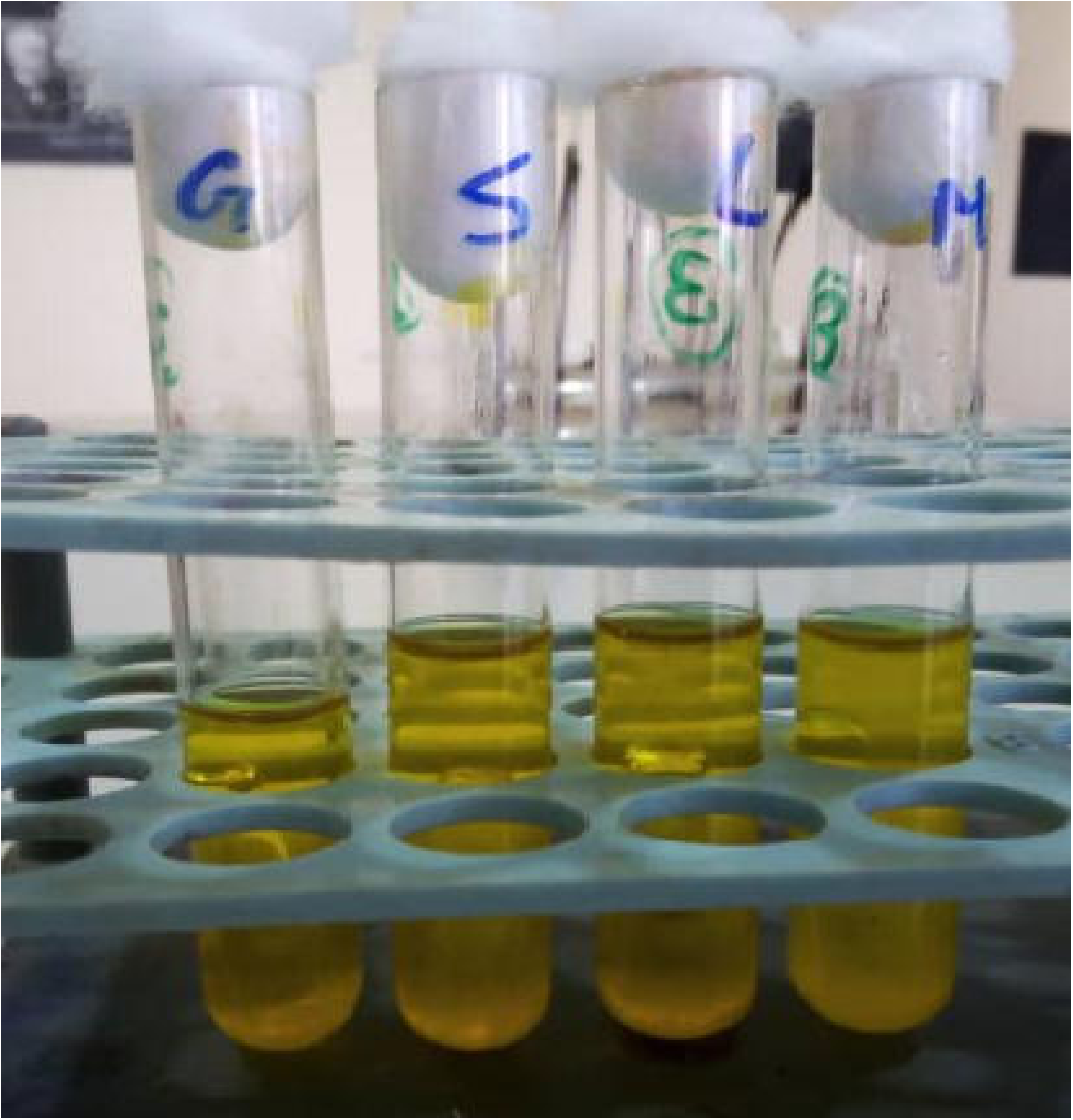
Sugar fermentation tests (Glucose, Sucrose, Lactose, Mannitol)

**Figure 5:**
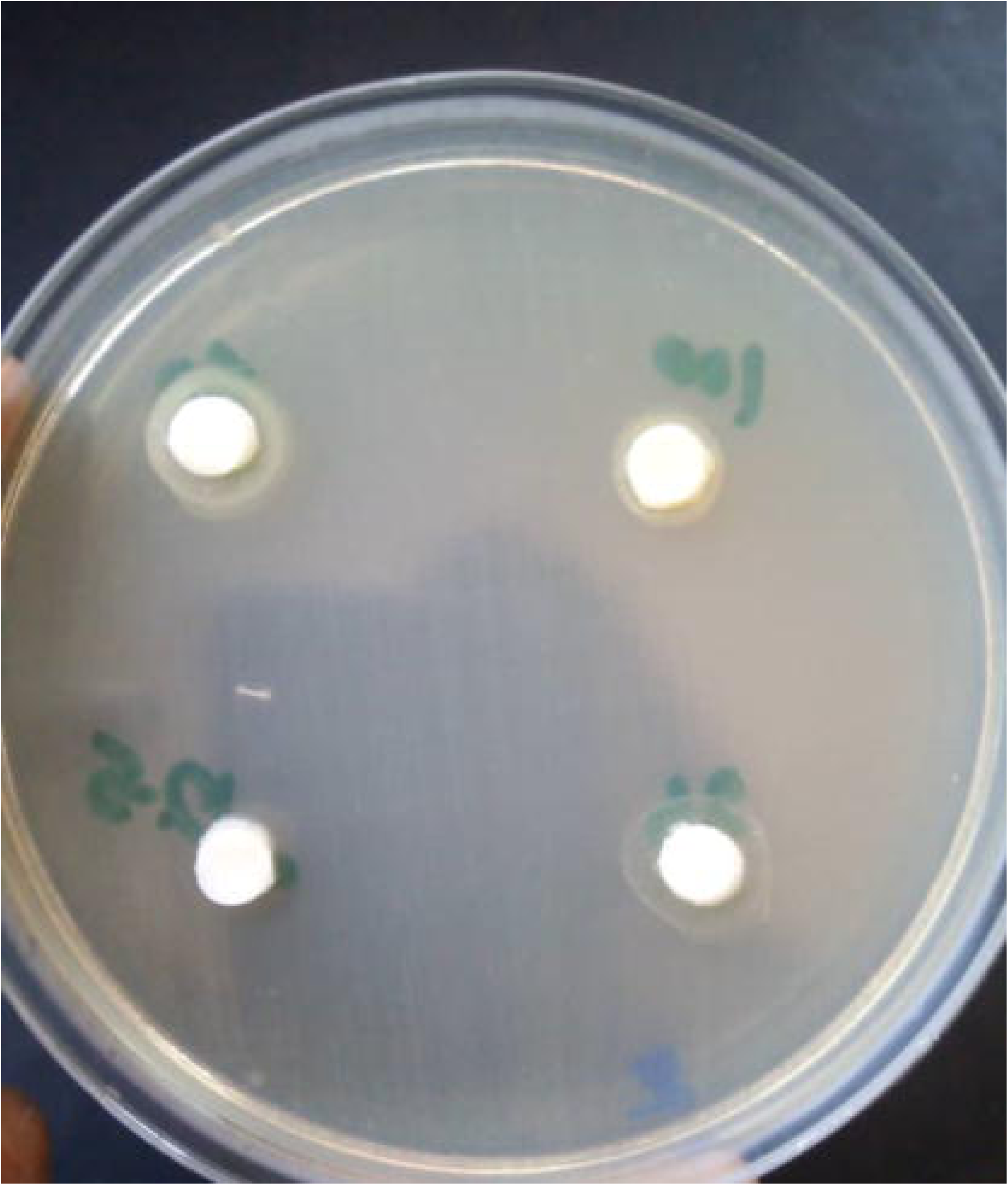
Antibacterial activity of *C.myrrha* oil against *Lactobacillus spp*. Antibiotics were tested against *S.mutans* and *Lactobacillus ssp*. Results showed in “**Table 3**”. The inhibition zone of Ampicillin using Disc diffusion method against *S.mutans* and *Lactobacillus ssp*. were found to be 17.7 mm ± 2.5 respectively. While *Lactobacillus ssp*. was resistant to Ampicillin. Vancomycin had inhibition zones of 23.7 mm ± 0.2 followed by Ciprofloxacin with inhibition zone of 18.3 mm ±1.5 against *Lactobacillus ssp*.

### Determination of Minimum Bactericidal Concentration (MBC)

With the three concentrations of *C.myrrha* oil at 12.5, 6.25 and 3.125 mg/ml no growth was recorded across the three isolates. However, growth of each of three isolates was observed at *C.myrrha* oil concentration 1.56 mg/ml “**Table 4**” The concentration of *C.myrrha* oil 3.125 mg/ml was considered as the (MBC).

**Table 3:** Antibacterial activity of antibiotics against bacterial strains.

|  | Inhibition zones measured in mm |  |  |  |
| --- | --- | --- | --- | --- |
|  | Mean | Std. | Min | Max |
| **Bacteria/Antibiotics** |  |  |  |  |
| *Streptococcus mutans* (n=3) |  |  |  |  |
| Ampicillin | 17.7 | 2.5 | 15.0 | 20.0 |
| *Lactobacillus ssp.* (n=3) |  |  |  |  |
| Ampicillin | 0.0 | 0.0 | 0.0 | 0.0 |
| Vancomycin | 23.7 | 1.2 | 23.0 | 25.0 |
| Ciprofloxacin | 18.3 | 1.5 | 17.0 | 20.0 |

**Table 4:**
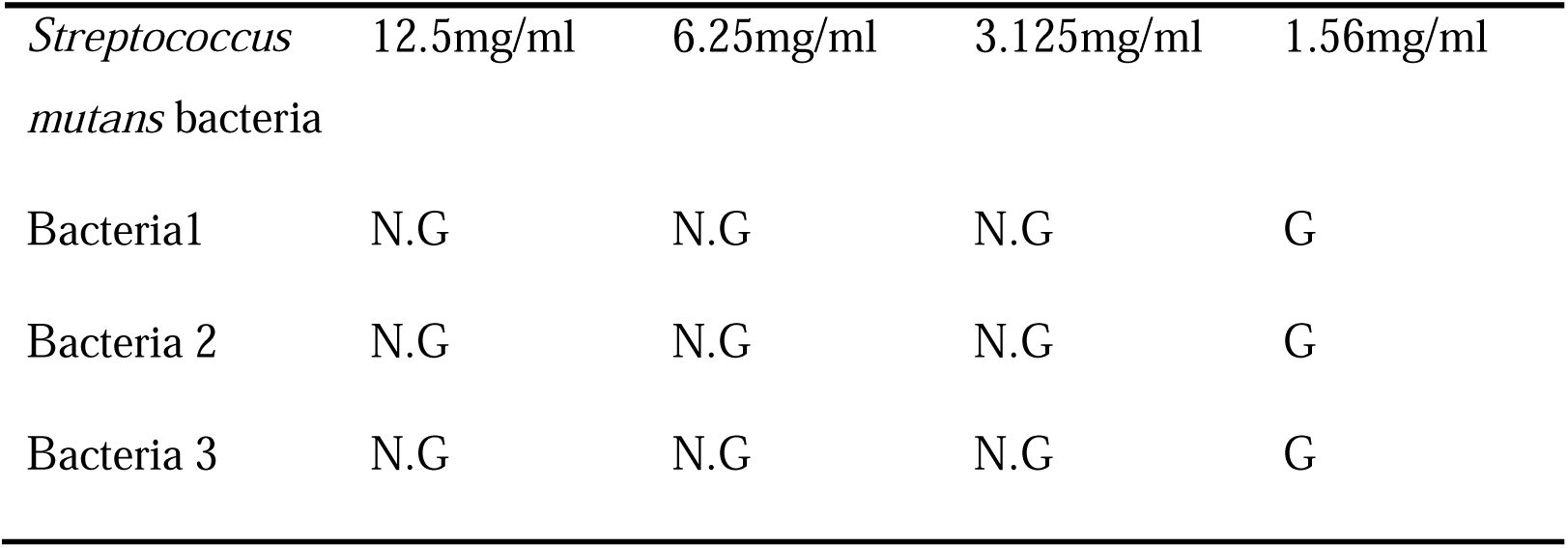
Growth of bacteria with four concentrations, (N.G: no growth; G: growth of bacteria).

### Gas Chromatography Mass Spectrometry

The results of GC/MS revealed 48 compounds after comparing retention index and mass fragmentation patents of the oil component with those available in the library, the National Institute of Standards and Technology (NIST). The highest percentage compounds were; Benzofuran 29.13% with retention time (R.T) 14.922 followed by Cyclohexane 19.88% with R.T 12.986 and 1,3-Diphenyle-1,2-butanediol 15,17% with R.T 17.134 “**Tables 5a and 5b**”

**Table 5a:** Component of oil as analysed by GC-MS.

| SN | Name | Retention Time | Formula | Molecular weight | Retention index | Area% |
| --- | --- | --- | --- | --- | --- | --- |
| 1 | Bicyclo[3.1.0]hex-2-ene, 2-methyl-5-(1-methylethyl)- | 4.125 | C<sub>10</sub>H<sub>16</sub> | 136 | 902 | 0.02 |
| 2 | .alpha.-Pinene | 4.244 | C<sub>10</sub>H<sub>16</sub> | 136 | 948 | 0.05 |
| 3 | Camphene | 4.495 | C<sub>10</sub>H<sub>16</sub> | 136 | 943 | 0.08 |
| 4 | .beta.-Pinene | 4.973 | C<sub>10</sub>H<sub>16</sub> | 136 | 943 | 0.02 |
| 5 | D-Limonene | 5.916 | C<sub>10</sub>H<sub>16</sub> | 136 | 1018 | 0.02 |
| 6 | Bicyclo[2.2.1]heptane, 2-methoxy-1,7,7-trimethyl- | 7.546 | C<sub>11</sub>H<sub>20</sub>O | 168 | 1087 | 0.02 |
| 7 | (+)-2-Bornanone | 8.237 | C<sub>10</sub>H<sub>16</sub>O | 152 | 1121 | 0.01 |
| 8 | 1,5-Cyclooctadiene, 3,4-dimethyl- | 10.247 | C<sub>10</sub>H<sub>16</sub> | 136 | 1046 | 0.03 |
| 9 | Acetic acid, 1,7,7-trimethyl-bicyclo[2.2.1]hept-2-yl ester | 10.97 | C<sub>12</sub>H<sub>20</sub>O<sub>2</sub> | 196 | 1277 | 0.11 |
| 10 | Cyclohexene, 4-ethenyl-4-methyl-3-(1-methylethenyl)-1-(1-methylethyl)-, (3R-trans)- | 11.929 | C<sub>15</sub>H<sub>24</sub> | 204 | 1377 | 4.56 |
| 11 | .alpha.-Cubebene | 12.154 | C<sub>15</sub>H<sub>24</sub> | 204 | 1344 | 0.24 |
| 12 | .gamma.-Muurolene | 12.42 | C<sub>15</sub>H<sub>24</sub> | 204 | 1435 | 0.04 |
| 13 | .alpha.-ylangene | 12.587 | C<sub>15</sub>H<sub>24</sub> | 204 | 1221 | 0.09 |
| 14 | Copaene | 12.67 | C<sub>15</sub>H<sub>24</sub> | 204 | 1221 | 0.37 |
| 15 | (-)-.beta.-Bourbonene | 12.851 | C<sub>15</sub>H<sub>24</sub> | 204 | 1339 | 1.78 |
| 16 | Cyclohexane, 1-ethenyl-1-methyl-2,4-bis(1-methylethenyl)-, [1S-(1.alpha.,2.beta.,4.beta.)]- | 12.986 | C<sub>15</sub>H<sub>24</sub> | 204 | 1398 | 19.88 |
| 17 | Caryophyllene | 13.5 | C<sub>15</sub>H<sub>24</sub> | 204 | 1494 | 1.76 |
| 18 | .gamma.-Elemene | 13.571 | C<sub>15</sub>H<sub>24</sub> | 204 | 1431 | 0.14 |
| 19 | .beta.-copaene | 13.671 | C<sub>15</sub>H<sub>24</sub> | 204 | 1216 | 0.25 |
| 20 | 1,5-Cyclodecadiene, 1,5-dimethyl-8-(1-methylethylidene)-, (E,E)- | 13.721 | C<sub>15</sub>H<sub>24</sub> | 204 | 1603 | 6.76 |
| 21 | .beta.-ylangene | 13.944 | C<sub>15</sub>H<sub>24</sub> | 204 | 1216 | 0.23 |
| 22 | Humulene | 14.122 | C<sub>15</sub>H<sub>24</sub> | 204 | 1579 | 0.58 |
| 23 | Alloaromadendrene | 14.253 | C<sub>15</sub>H<sub>24</sub> | 204 | 1386 | 0.15 |
| 24 | 1,6-Cyclodecadiene, 1-methyl-5-methylene-8-(1-methylethyl)-, [S-(E,E)]- | 14.353 | C<sub>15</sub>H<sub>24</sub> | 204 | 1515 | 0.05 |
| 25 | 4,5-di-epi-aristolochene | 14.401 | C<sub>15</sub>H<sub>24</sub> | 204 | 1474 | 0.09 |
| 26 | Guaia-1(10),11-diene | 14.508 | C<sub>15</sub>H<sub>24</sub> | 204 | 1490 | 1 |
| 27 | 1H-Cyclopenta[1,3]cyclopropa[1,2]benzene, octahydro-7-methyl-3-methylene-4-(1-methylethyl)-, [3aS-(3a.alpha.,3b.beta.,4.beta., | 14.611 | C<sub>15</sub>H<sub>24</sub> | 204 | 1339 | 1.43 |
| 28 | Naphthalene, decahydro-4a-methyl-1-methylene-7-(1-methylethenyl)-, [4aR-(4a.alpha.,7.alpha.,8a.beta.)]- | 14.715 | C<sub>15</sub>H<sub>24</sub> | 204 | 1469 | 2.48 |
| 29 | Benzofuran, 6-ethenyl-4,5,6,7-tetrahydro-3,6-dimethyl-5-isopropenyl-, trans- | 14.922 | C<sub>15</sub>H<sub>20</sub>O | 216 | 1532 | 29.13 |

**Table 5b:** Component of oil as analysed by GC-MS.

| S<br>N | Name | Retention<br>Time | Formula | Molecular<br>weight | Retention<br>index | Area% |
| --- | --- | --- | --- | --- | --- | --- |
| 30 | 1,4-Methano-1H-indene, octahydro-4-methyl-8-methylene-7-(1-methylethyl)-, [1S-(1.alpha.,3a.beta.,4.alpha.,7.alpha.,7a.beta.)]- | 15.057 | C<sub>15</sub>H<sub>24</sub> | 204 | 1339 | 0.27 |
| 31 | Naphthalene, 1,2,3,4,4a,5,6,8a-octahydro-7-methyl-4-methylene-1-(1-methylethyl)-, (1.alpha.,4a.beta.,8a.alpha.)- | 15.187 | C<sub>15</sub>H<sub>24</sub> | 204 | 1435 | 0.57 |
| 32 | Naphthalene, 1,2,3,5,6,8a-hexahydro-4,7-dimethyl-1-(1-methylethyl)-, (1S-cis)- | 15.324 | C<sub>15</sub>H<sub>24</sub> | 204 | 1469 | 0.57 |
| 33 | Naphthalene, 1,2,3,5,6,7,8,8a-octahydro-1,8a-dimethyl-7-(1-methylethenyl)-, [1S-(1.alpha.,7.alpha.,8a.alpha.)]- | 15.56 | C<sub>15</sub>H<sub>24</sub> | 204 | 1474 | 0.54 |
| 34 | Selina-3,7(11)-diene | 15.67 | C<sub>15</sub>H<sub>24</sub> | 204 | 1507 | 0.5 |
| 35 | Azulene, 1,2,3,3a,4,5,6,7-octahydro-1,4-dimethyl-7-(1-methylethenyl)-, [1R-(1.alpha.,3a.beta.,4.alpha.,7.beta.)]- | 15.947 | C<sub>15</sub>H<sub>24</sub> | 204 | 1461 | 1.31 |
| 36 | Carbamic acid, p-tolyl ester | 16.318 | C<sub>9</sub>H<sub>11</sub>NO<sub>2</sub> | 165 | 1372 | 0.37 |
| 37 | Caryophyllene oxide | 16.415 | C<sub>15</sub>H<sub>24</sub>O | 220 | 1507 | 0.11 |
| 38 | Cyclohexanone, 5-ethenyl-5-methyl-4-(1-methylethenyl)-2-(1-methylethylidene)-, cis- | 16.741 | C<sub>15</sub>H<sub>22</sub>O | 218 | 1602 | 0.13 |
| 39 | cis-Z-.alpha.-Bisabolene epoxide | 16.855 | C<sub>15</sub>H<sub>24</sub>O | 220 | 1531 | 0.07 |
| 40 | 1,3-Diphenyl-1,2-butanediol | 17.134 | C<sub>16</sub>H<sub>18</sub>O<sub>2</sub> | 242 | 2025 | 15.17 |
| 41 | 4,4'-Dimethyl-2,2'-dimethylenebicyclohexyl-3,3'-diene | 17.244 | C<sub>16</sub>H<sub>22</sub> | 214 | 1618 | 5.88 |
| 42 | Cyclohexanemethanol, 4-ethenyl-.alpha.,.alpha.,4-trimethyl-3-(1-methylethenyl)-, [1R-(1.alpha.,3.alpha.,4.beta.)]- | 17.835 | C<sub>15</sub>H<sub>26</sub>O | 222 | 1522 | 0.62 |
| 43 | Spiro[2.4]heptane, 1,2,4,5-tetramethyl-6-methylene- | 18.095 | C<sub>12</sub>H<sub>22</sub> | 164 | 1065 | 0.44 |
| 44 | Cyclohexene, 4-(1,5-dimethyl-1,4-hexadienyl)-1-methyl- | 18.294 | C<sub>15</sub>H<sub>24</sub> | 204 | 1518 | 0.37 |
| 45 | Bicyclo[3.1.1]hept-2-ene-2-ethanol, 6,6-dimethyl-, (1R)- | 18.56 | C<sub>11</sub>H<sub>18</sub>O | 166 | 1290 | 0.83 |
| 46 | Androsta-3,5-dien-7-one | 19.902 | C<sub>19</sub>H<sub>26</sub>O | 270 | 1933 | 0.4 |
| 47 | Acetic acid, 6-(1-hydroxymethyl-vinyl)-4,8a-dimethyl-3-oxo-1,2,3,5,6,7,8,8a-octahydronaphthalen-2-yl ester | 20.871 | C<sub>17</sub>H<sub>24</sub>O<sub>4</sub> | 292 | 2244 | 0.27 |
| 48 | .alpha.-Bulnesene | 21.362 | C<sub>15</sub>H<sub>24</sub> | 204 | 1490 | 0.21 |

## Discussion

Medicinal plant has been used more commonly as medications due to it is availability and safety and as the resistance to antibiotics drugs is successively increase, According to the results, *C.myrrha* has been used in oral health to treat periodontitis and malodour [11,12]. *S.mutans* and *Lactobacillus ssp*. bacteria responsible in formation and progression of dental caries were tested with *C.myrrha* oil using two different methods [5]. In this study *C.myrrha* oil was effective at all concentrations against *S.mutans* bacteria with MBC 3.125mg/ml while *Lactobacillus ssp*. MBC was considered to be 25mg/ml. The highest activity of oil against *S.mutans* and *Lactobacillus rhamnosus* using disc diffusion method were found to be 18.34 mm±0.26 and 11.72 mm±0.85 respectively. Higher results were recorded in a comparative study testing two different formula of Myrrh mouth wash (containing 65 ml of Myrrh extract for each 100ml) with another ingredient against *Streptococcus mutans* isolates. Their findings for the two formulas were 32.367 mm ± 0.262 and 22.367 mm ± 0.102 respectively [14]. In this study, the result of antimicrobial activity of Ampicillin antibiotic against *S.mutans* using disc method was comparable to result of myrrh against same bacteria .Myrrh can be used as alternative and effective antibacterial agent under condition that further investigations are conducted. Furthermore, *Lactobacillus ssp*. was completely resistant to Ampicillin while Myrrh was effective at the three concentrations of respectively 100,50, and 25 mg/ml. Vancomycin was more effective than Ciprofloxacin with inhibition zones of respectively 23.7 mm ±1.2 and 18.3 mm ±1.5.

Sesquiterpenes and furanotype, are major constituents of *C.myrrha*, have antimicrobial activity leading them to have therapeutic effect [10,14,20]. The highest percentage compound in our result is Benzofuran 29.13%, which have antimicrobial activity, this results in line with a study which revealed that benzofuran constitutes 26.63% of *C.myrrha* oil analysed by GC-MS technique [20], and higher than another study which indicated that Curzerene (Benzofuran) was 11.9% [9].

Caryophyllene C_15_H_24_ considered as anti-tumor, antibacterial, anti-inflammatory represents 1.76% of constituents higher than a study which revealed 0.29% [20]. Other important components of myrrh oil in our study, measured through R.T 12.986 and R.T 17.134., were Cyclohexane (19.88 %) and 1,3-Diphenyle-1,2-butanediol (15.17%). Myrrh oil was effective on both *S.mutans* and *Lactobacillus ssp*. with a higher activity on *S.mutans*. The two bacteria are involved in dental caries; hence, Myrrh oil is a potential antibacterial product that needs to be recognized as such by further scientific studies and may be good for oral health to control caries as a public health challenge.

## Conclusion

The results obtained support some of the traditional uses of Myrrh and may offer potential leads to new active natural products. Further researches are needed to identify the active principles and development new drugs against various diseases.

## Acknowledgments

Special thanks to Afraa M.Alhaj, Department of Microbiology, Faculty of Medical Laboratory Sciences, University of Medical Sciences and Technology. for facilitating laboratory experiments. Special thanks to the faculty of Medical Laboratory Sciences, University of Medical Sciences and Technology in which I process this study, for their support. Greatest thanks to Khartoum Dental Teaching hospital for help and support. Also, I would like to thanks Zawahir Abu Elbashar, Department of Diary Science and Technology, Sudan University of Science and Technology. for providing the research with lactobacillus bacteria

